# Signaling from the RNA sensor RIG-I is regulated by ufmylation

**DOI:** 10.1101/2021.10.26.465929

**Authors:** Daltry L. Snider, Moonhee Park, Kristen A. Murphy, Dia C. Beachboard, Stacy M. Horner

## Abstract

The RNA binding protein RIG-I is a key initiator of the antiviral innate immune response. The signaling that mediates the antiviral response downstream of RIG-I is transduced through the adaptor protein MAVS and results in the induction of type I and III interferons (IFN). This signal transduction occurs at endoplasmic reticulum (ER)-mitochondrial contact sites, to which RIG-I and other signaling proteins are recruited following their activation. RIG-I signaling is highly regulated to prevent aberrant activation of this pathway and dysregulated induction of IFN. Previously, we identified UFL1, the E3 ligase of the ubiquitin-like modifier conjugation system called ufmylation, UFL1, as one of the proteins recruited to membranes at ER-mitochondrial contact sites in response to RIG-I activation. Here, we show that UFL1, as well as the process of ufmylation, promote IFN induction in response to RIG-I activation. We find that following RNA virus infection, UFL1 is recruited to the membrane targeting protein 14-3-3ε, and that this complex is then recruited to activated RIG-I to promote downstream innate immune signaling. Importantly, we found that 14-3-3ε has an increase in UFM1-conjugation following RIG-I activation. Additionally, loss of cellular ufmylation prevents the interaction of 14-3-3ε with RIG-I, which abrogates the interaction of RIG-I with MAVS and thus downstream signal transduction that induces IFN. Our results define ufmylation as an integral regulatory component of the RIG-I signaling pathway and as a post-translational control for IFN induction.

**Significance:** The viral RNA sensor RIG-I initiates the antiviral innate immune response by activating a signaling cascade that induces interferon. Activation of the RIG-I signaling pathway is highly regulated to quickly mount a protective immune response while preventing dysregulation that can lead to excessive inflammation or autoimmune disorders. Here, we characterize one such mechanism of regulation. We describe that UFL1, an E3 ligase for the ubiquitin-like modifier conjugation system called ufmylation, is important to promote RIG-I signaling. Using molecular approaches, we show that ufmylation promotes RIG-I interaction with the membrane targeting protein 14-3-3ε. As such, ufmylation positively regulates RIG-I recruitment to its signaling adaptor proteins MAVS for induction of interferon in response to RNA virus infection.

## Introduction

Detection of RNA virus infection is initiated by cellular sensors such as RIG-I. RIG-I is a pattern recognition receptor that detects unique features of viral RNA that are generally absent in cellular RNA, referred to as pathogen-associated molecular patterns (PAMPs) (1). Sensing of viral RNA PAMPs triggers RIG-I activation and induces a downstream signaling cascade that ultimately results in transcriptional induction of type I and type III interferons (IFN) and the antiviral response (2, 3). The RIG-I signaling cascade is carefully regulated by multiple mechanisms, including post-translational modifications that influence specific protein-protein interactions that can result in changes in protein localization to mediated signaling (3, 4). For example, following sensing of RNA PAMPs, RIG-I undergoes K63-linked polyubiquitination in order to transition to its fully active conformation, which promotes its interaction with the molecular trafficking protein 14-3-3ε (5–8). 14-3-3ε facilitates the recruitment of activated RIG-I from the cytosol to intracellular membranes where it interacts with MAVS (7, 9, 10), which assembles other RIG-I pathway members to transduce the signals that induce IFN (7, 11). Importantly, many RNA viruses, including influenza A virus and some flaviviruses (dengue virus, Zika virus, and West Nile virus), prevent the interaction of RIG-I with 14-3-3ε to limit IFN induction and evade the antiviral response (9, 10, 12).

In addition to RIG-I, a number of signaling proteins must be recruited to MAVS in order to propagate downstream IFN induction. Previously, we identified proteins that move to MAVS signaling sites at mitochondrial-associated endoplasmic reticulum (ER) membranes (MAM) during RNA virus infection (13, 14). These proteins likely aid in spatial organization of RIG-I pathway proteins during viral infection and include the GTPase RAB1B, which plays a role in recruiting TRAF3 to MAVS (15). In addition to RAB1B, we identified other proteins recruited to the MAM upon RIG-I signaling activation, one of which was UFL1 (referred to in our previous publication as KIAA0776) (14). UFL1 is an E3 ligase for UFM1, which is a ubiquitin-like modification of 85 amino acids. The process of ufmylation conjugates UFM1 covalently to lysine residues of target proteins through a process called ufmylation, which is similar to ubiquitination in that it also uses an E1, E2, and E3 ligase conjugation system (UBA5, UFC1, and UFL1; see Figure 2D). UFM1 is removed by the UFSP2 protease (16–20). The consequence of UFM1 addition to proteins is not fully understood, but the literature supports the idea that it can promote protein protein interactions to regulate a number of biological processes (21–31). Here, we uncover a role for ufmylation in RIG-I activation. We found that the cellular proteins that catalyze ufmylation all promote RIG-I-mediated induction of IFN. Interestingly, we found that following RNA virus infection UFL1 interacts with both RIG-I and the molecular trafficking protein 14-3-3ε, and that 14-3-3ε undergoes ufmylation. Further, similar to RIG-I, UFL1 is recruited to intracellular membranes following RNA virus infection. Importantly, loss of ufmylation prevents the interaction of 14-3-3ε with RIG-I, which results in decreased MAVS activation and IFN induction in response to RNA virus infection. Thus, ufmylation can regulate RIG-I activation and downstream signaling of the intracellular innate immune system.

## Results

### The ufmylation activity of UFL1 promotes RIG-I signaling

Having found that the E3 ligase of ufmylation UFL1 is recruited to MAVS signaling sites at the MAM in response to RIG-I signaling (14), we wanted to determine if UFL1 regulates RIG-I signaling. To test this, we measured induction of the IFN-β promoter following UFL1 overexpression using an IFN-β promoter luciferase reporter assay (32) and found that UFL1 increased activation of the IFN-β promoter, similar to that of RIG-I expression, in a dose-dependent fashion in response to infection with Sendai virus (SenV) (Figure 1A). SenV is a murine paramyxovirus that specifically activates RIG-I (33). In support of UFL1 enhancing RIG-I signaling specifically, exogenous expression of UFL1 also increased IFN-β promoter activity in response to transfection of 293T cells with a known RIG-I immunostimulatory RNA from hepatitis C virus (PAMP; Figure S1A) (34). However, UFL1 overexpression in 293T cells did not lead to increased induction of IFN-stimulated genes (ISG), such as *ISG56* or *ISG15*, in response to exogenous IFN-β treatment, which bypasses RIG-I signaling and IFN induction, indicating that UFL1 primarily regulates IFN induction and not the IFN response (Figure S1B). Next, we depleted UFL1 by siRNA in two different cell types and measured SenV-induced activation of the RIG-I pathway. Depletion of UFL1 in primary neonatal human dermal fibroblasts (NHDFs) reduced the SenV-mediated induction of both *IFNB1* and *IFNL1* transcripts, as measured by RT-qPCR (Figure 1B), as well as production of IFN-β protein, as measured by an enzyme-linked immunosorbent assay (ELISA) (Figure 1C). Depletion of UFL1 in 293T cells resulted in decreased phosphorylation of IRF3, a transcription factor for both type I and III IFNs, while exogenous expression of an siRNA-resistant UFL1 restored SenV-mediated IRF3 phosphorylation (Figure 1D).

**Figure 1.**
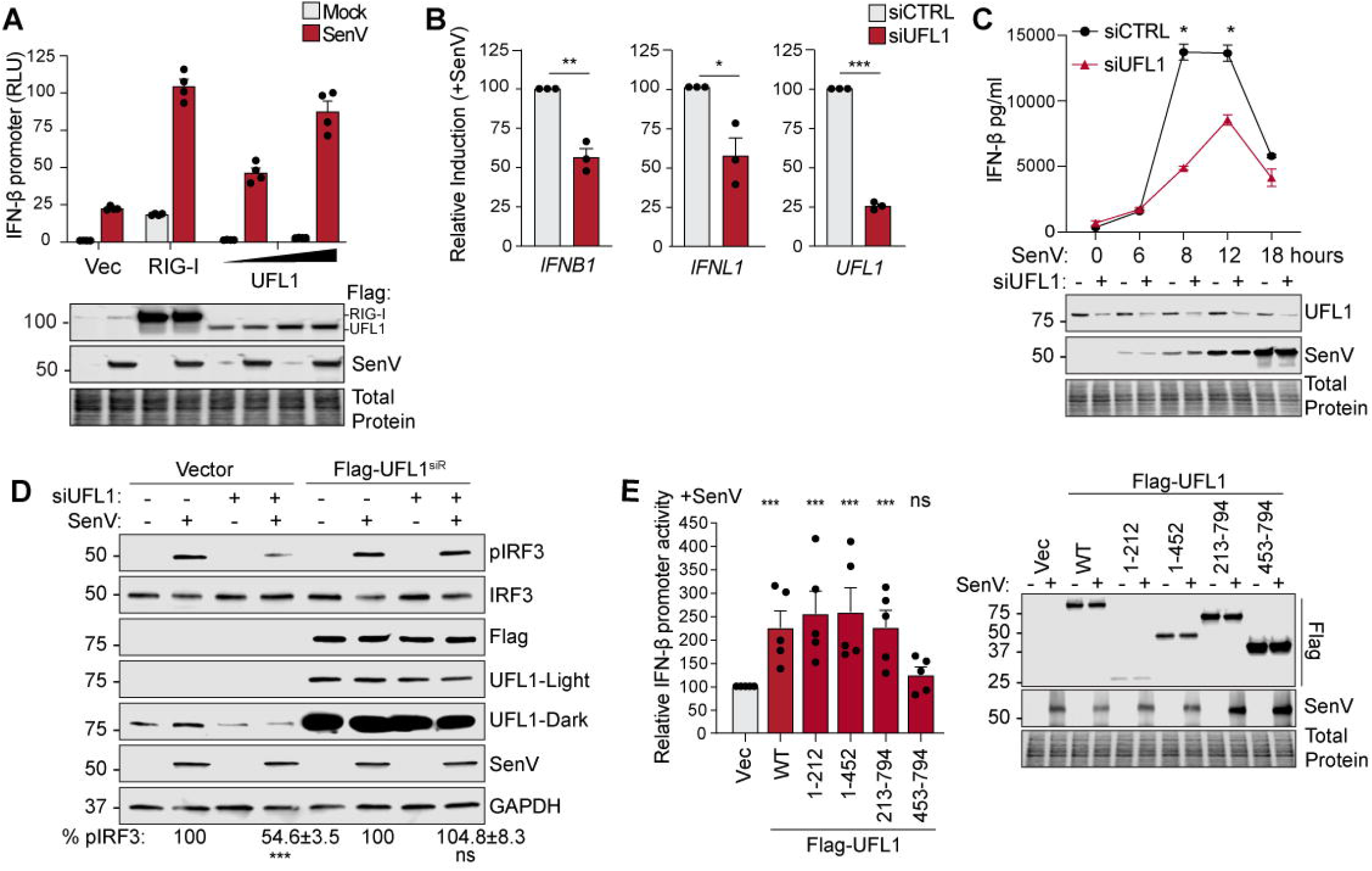
The ufmylation activity of UFL1 promotes RIG-I signaling. A) IFN-β-promoter reporter luciferase expression (rel. to CMV-*Renilla*) from 293T cells expressing vector, Flag-UFL1, or Flag-RIG-I, followed by mock or SenV infection (18 h). B) RT-qPCR analysis (rel. to *18S*) of RNA extracted from primary neonatal human dermal fibroblasts (NHDFs) transfected with either siCTRL or siUFL1 followed by mock or SenV infection (8 h). C) ELISA for IFN-β of supernatants harvested from NHDFs transfected with siCTRL or siUFL1 and infected with SenV for the indicated times. D) Immunoblot analysis of p-IRF3 following siRNA transfection along with expression of vector or Flag-UFL1^SiR^, which has point mutations in the siRNA seed sequence. Quantification of p-IRF3/Tubulin is shown on underneath. E) Relative IFN-β-promoter reporter luciferase expression (rel. to CMV-*Renilla*) from 293T cells expressing indicated constructs followed by mock or SenV infection (12-18 h), with results graphed as relative SenV fold change for each. For A) mean -/+ SD, n=3 technical replicates and representative of n=3 independent experiments. For all others, mean -/+ SEM, n=3 or n=5 (1E) biological replicates. *p ≤ 0.05, **p ≤ 0.01, and ***p ≤ 0.001 determined by one-way ANOVA followed by Šidák’s multiple comparisons test (C), Student’s t-test (B, D), or one-way ANOVA followed by Dunnett’s multiple comparisons test (E).

To define the domains of UFL1 that regulate RIG-I signaling, we expressed a series of previously described UFL1 truncation mutants and measured SenV-mediated activation of the IFN-β promoter in a luciferase reporter assay (16). The ability of UFL1 to transfer UFM1 to a target protein has been suggested to require the first 212 amino acids of the protein, as this domain interacts with the E2 ligase for ufmylation, UFC1 (16). The wild-type (WT) UFL1 (aa 1-794), as well as the C-terminal deleted mutants of UFL1, aa 1-212 and aa 1-452, which all have reported ufmylation activity (16), stimulated SenV-medicated induction of the IFN-β promoter (Figure 1E). Interestingly, the N-terminal deleted mutant of UFL1 aa 213-794, which does not have reported ufmylation activity, also induced signaling, while the N-terminal deleted UFL1 mutant aa 453-794 did not (Figure 1E). To measure UFL1 conjugation activity in our hands, we quantified the formation of higher molecular weight UFM1-conjugates by immunoblotting. The 10 kDa UFM1 protein is known to be conjugated to a number of proteins, including the highly abundant RPL26 (21, 28). Indeed, our analysis of the higher molecular weight UFM1-conjugates induced by these UFL1 constructs revealed differential abundance of these UFM1-conjugates. Specifically, UFL1 WT, aa 1-212, aa 1-452, and aa 213-794 of UFL1 all retain full ufmylation activity, while aa 453-794 of UFL1 does not (Figure S1C). Thus, taken together, this reveals that the ufmylation activity of UFL1 is required to promote RIG-I signaling that results in induction of IFN.

### The ufmylation machinery proteins positively regulate RIG-I signaling

Having determined that the ufmylation activity of UFL1 is important for its role in RIG-I signaling, we hypothesized that UFM1 and the proteins required for UFM1 conjugation would also be required to promote this signaling. Similar to our results with UFL1, overexpression of UFM1 increased SenV-mediated activation of the IFN-β promoter, resembling the magnitude of induction seen with RIG-I, in a dose-dependent fashion (Figure 2A). Conversely, the activation of the IFN-β promoter in response to SenV was significantly abrogated in 293T cells in which UFM1 was deleted by CRISPR/Cas9, as compared to WT 293T cells (Figure 2B). Importantly, this signaling was restored upon exogenous expression of UFM1 (Figure 2B). The absence of UFM1 expression also prevented the induction of IFN-β protein in response to SenV infection, as measured by ELISA (Figure 2C). The process of ufmylation has 5 steps (Figure 2D). First, UFM1 is processed to expose the terminal glycine residue. Then, this mature UFM1 is added to the target protein by the actions of UBA5, which acts as an E1 ligase for UFM1; UFC1, the E2 ligase; and UFL1, the E3 ligase (19). Finally, the UFSP2 protease removes UFM1, which enables recycling of mature UFM1 (18). We found that exogenous expression of each of the proteins involved in UFM1 conjugation, including the UFSP2 protease, positively regulated SenV-mediated induction of the IFN-β promoter (Figure 2E). Conversely, the induction of *IFNB1* by SenV was abrogated in 293T cells in which UFSP2 was deleted by CRISPR/Cas9, similar to that seen upon UFM1 KO. Interestingly, we observed an increase in UFM1-retention by an ufmylated protein and a decrease in free UFM1 in the UFSP2 KO cells via anti-UFM1 immunoblot, suggesting UFSP2 may serve to primarily to increase the pool of available mature UFM1 for conjugation (Figure 2F). These results reveal that the proteins that catalyze ufmylation and the UFM1 modification itself promote RIG-I signaling.

**Figure 2.**
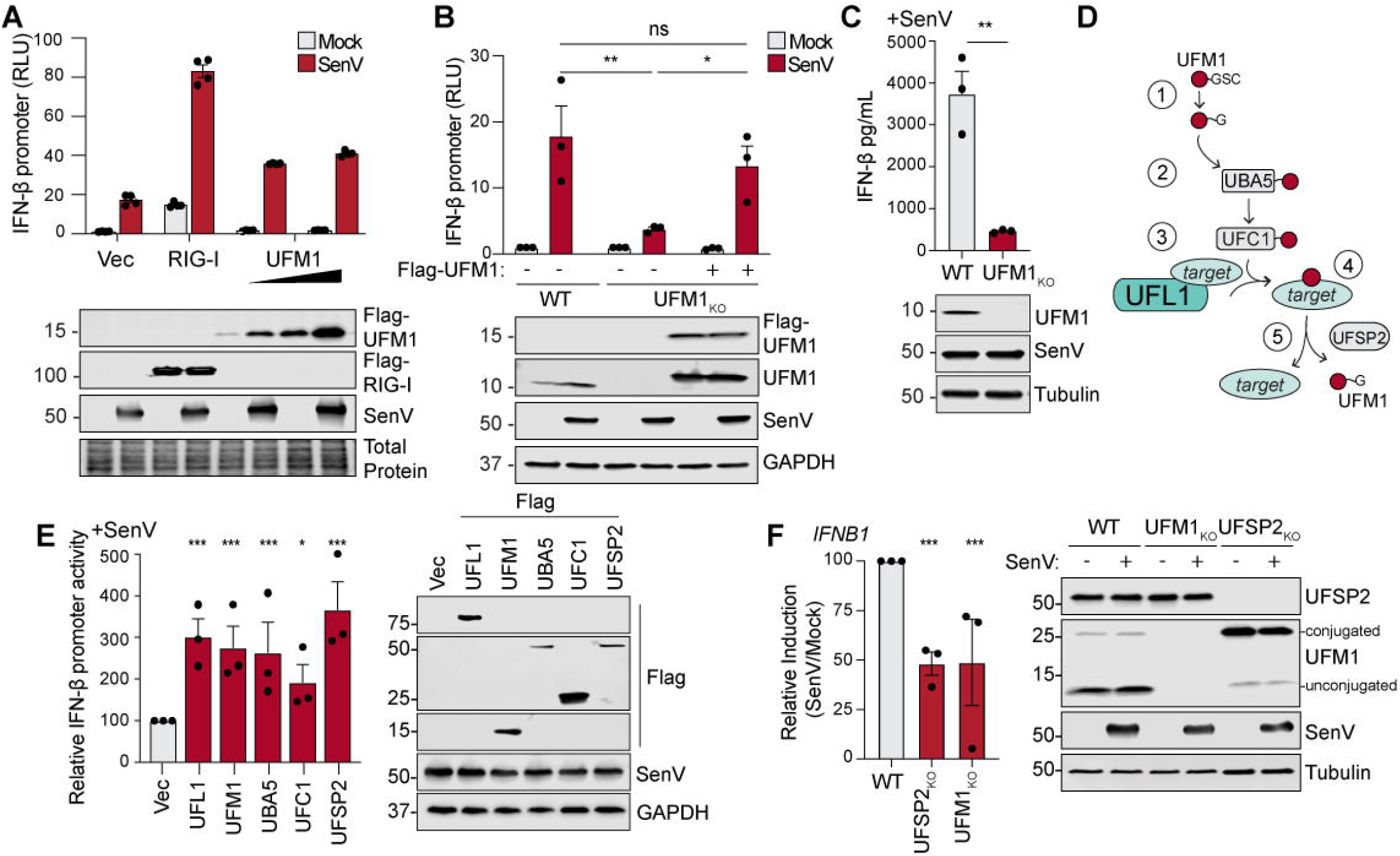
The ufmylation machinery proteins positively regulate RIG-I signaling. A) IFN-β-promoter reporter luciferase expression (rel. to CMV-*Renilla*) from 293T cells expressing vector, Flag-UFM1, or Flag-RIG-I followed by mock or SenV infection (18 h) or in B) WT or CRISPR/CAS9 UFM1 KO 293T cells transfected with vector (Vec) or Flag-UFM1 (for KO), followed by mock or SenV infection (18 h). C) ELISA for IFN-β of supernatants harvested from WT or CRISPR/CAS9 UFM1 KO 293T cells that were SenV infected (18 h). D) Diagram of UFM1 conjugation. E) Relative IFN-β-promoter reporter luciferase expression (rel. to CMV-*Renilla*) from 293T cells expressing indicated constructs followed by mock or SenV infection (18 h), with results graphed as relative SenV fold change for each. F) RT-qPCR analysis (rel. to *GAPDH*) of RNA extracted from 293T WT, UFM1 KO or UFSP2 KO with either mock or SenV infection (18 h). For A) mean -/+ SD, n=3 technical replicates and representative of n=3 independent experiments. For all others, mean -/+ SEM, n=3 biological replicates. *p ≤ 0.05, **p ≤ 0.01, and ***p ≤ 0.001 determined by two-way ANOVA followed by Tukey’s multiple comparisons test (A-B), Student’s t-test (C), or one-way ANOVA followed by Dunnett’s multiple comparisons test (E, F).

### UFM1 is required for the RIG-I-driven transcriptional response

After establishing that ufmylation promotes RIG-I activation, and in turn IFN expression, we next broadly measured the impact of ufmylation upon the transcriptional response to RIG-I signaling. Using RNA-sequencing, we analyzed gene expression in either WT or UFM1 KO 293T cells, following mock or SenV infection (Table S1.1; Table S1.2). Gene set enrichment analysis (Table S2.1; Table S2.2) of the transcripts significantly reduced (adjusted P<0.01) by UFM1 KO in the absence of viral infection revealed previously described pathways regulated by ufmylation such as cytosolic ribosomes, ribosome assembly, and hematopoiesis (Figure S2A; Table S2.1) (21, 28, 29, 35). Following viral infection, the top 10 gene categories negatively impacted by UFM1 KO, with a darker red color indicating more downregulation, were all related to the antiviral response, such as response to type I IFN and defense against virus (Figure 3A; Table S2.2). Indeed, of the top 50 most downregulated pathways impacted by UFM1 KO during infection, the majority were related to innate immune signaling or viral replication (Table S2.2), while upregulated gene categories were more diverse (Table S2.3; Table S2.4). Of the genes differentially expressed during UFM1 KO in response to SenV (adjusted P<0.01), the majority are downregulated (Figure 3B). Indeed, these downregulated genes included *IFNB1* and *IFNL1*, as well as other known ISGs (in red) (36) (Figure 3B; Figure 3C). These data are consistent with a model in which ufmylation-mediated regulation of IFN induction has broad consequences on genes induced by the IFN response.

**Figure 3.**
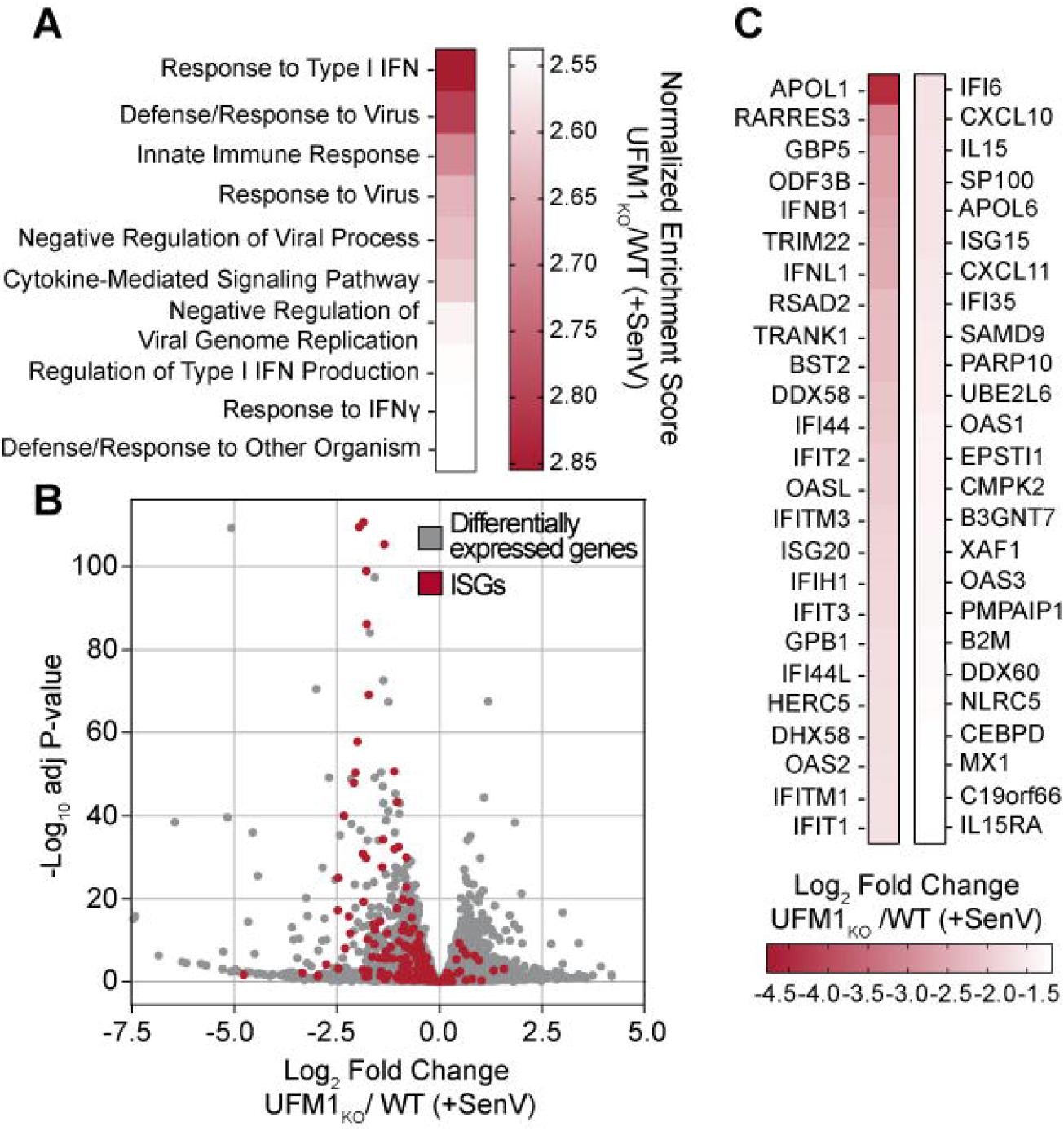
UFM1 is required for the RIG-I driven transcriptional response. RNA-seq analysis WT or UFM1 KO 293T cells following mock or SenV infection (18 h). A) Gene set enrichment analysis of negatively regulated differentially expressed genes in SenV-infected 293T cells represented by normalized enrichment score (UFM1 KO / WT). B) Volcano plot of differentially expressed genes (adj P<0.01) shown in grey, with ISGs shown in red, in SenV-infected 293T cells (UFM1 KO / WT). C) Heatmap of the effect of UFM1 KO on the fold change of the 50 most induced IFN and ISGs (UFM1 KO / WT) following SenV infection (adj P<0.01).

### UFL1 is recruited to intracellular membranes and interacts with 14-3-3ε and RIG-I during RNA virus infection

Following the binding of RIG-I to non-self RNA, it interacts with several host proteins to facilitate its activation, localization to the MAM, and interaction with MAVS. These proteins include the E3 ligases for K63-linked ubiquitin Riplet and TRIM25 (5, 6, 37), as well as the molecular trafficking protein 14-3-3ε. In particular, 14-3-3ε is required for RIG-I recruitment from the cytosol to MAVS signaling sites at intracellular membranes (5–7, 13); however, the mechanism underlying how 14-3-3ε selects RIG-I as cargo has yet to be elucidated. Using a subcellular membrane fractionation assay (38), we confirmed that UFL1 increases its association with intracellular membranes in response to SenV, similar to RIG-I (Figure 4A; compare fraction #1, which has Cox-I and no GAPDH, with fractions #6-8, which are enriched for the cytosolic protein GAPDH) (7, 12). This finding is consistent with our previous report that UFL1 is recruited to the MAM in response to either SenV or hepatitis C virus replication (14), suggesting that UFL1 recruitment occurs prior to MAVS activation, as MAVS is cleaved by the HCV NS3-NS4A protease (39–42). As the recruitment of RIG-I to intracellular membranes is known to require 14-3-3ε, and, as both UFL1 and UFM1 have been shown to interact with 14-3-3ε (16), we hypothesized that UFL1 may interact with 14-3-3ε to promote the IFN induction that we had observed in response to RNA virus infection. Thus, we first determined if the interaction of UFL1 with 14-3-3ε is increased in response to RIG-I activation by SenV by performing co-immunoprecipitation. We found that Myc-14-3-3ε did co-immunoprecipitate with Flag-UFL1, as reported previously (16), and that this interaction was increased by SenV (Figure 4B). Interestingly, the interaction of UFL1 with RIG-I also increased following SenV, both upon over-expression and at the level of the endogenous proteins (Figure 4C; Figure 4D). As RIG-I undergoes a series of modifications to become fully active (1, 4), we next used a panel of RIG-I mutants to define which stage of RIG-I activation promotes interaction with UFL1. These mutations prevent the distinct steps of RIG-I activation such as RIG-I binding to RNA (K888/907A), interacting with TRIM25 (T55I), or ubiquitination by Riplet and TRIM25 (K172/788R) (5, 43, 44). The interaction of UFL1 with RIG-I was significantly impaired by each of these mutations, suggesting that UFL1 regulates RIG-I function after it binds RNA and becomes ubiquitinated (Figure 4E). As this is the same step of activation at which 14-3-3ε binds to RIG-I to promote its translocation to intracellular membranes (7), we next tested the ability of UFL1 to promote RIG-I signaling in conjunction with the RIG-I regulatory factors Riplet and 14-3-3ε (6, 7). We found that co-expression of equal amounts of UFL1 with either Riplet or 14-3-3ε increased SenV-mediated activation of the IFN-β promoter above that seen by the individual proteins (Figure 4F). Intriguingly, the expression of UFL1 appears to be stabilized or enhances in the presence of Riplet and 14-3-3ε. These data together suggest that RNA virus infection increases the interaction of 14-3-3ε with UFL1, which then interacts with activated, K63-ubiquitinated RIG-I to promote downstream signaling.

**Figure 4.**
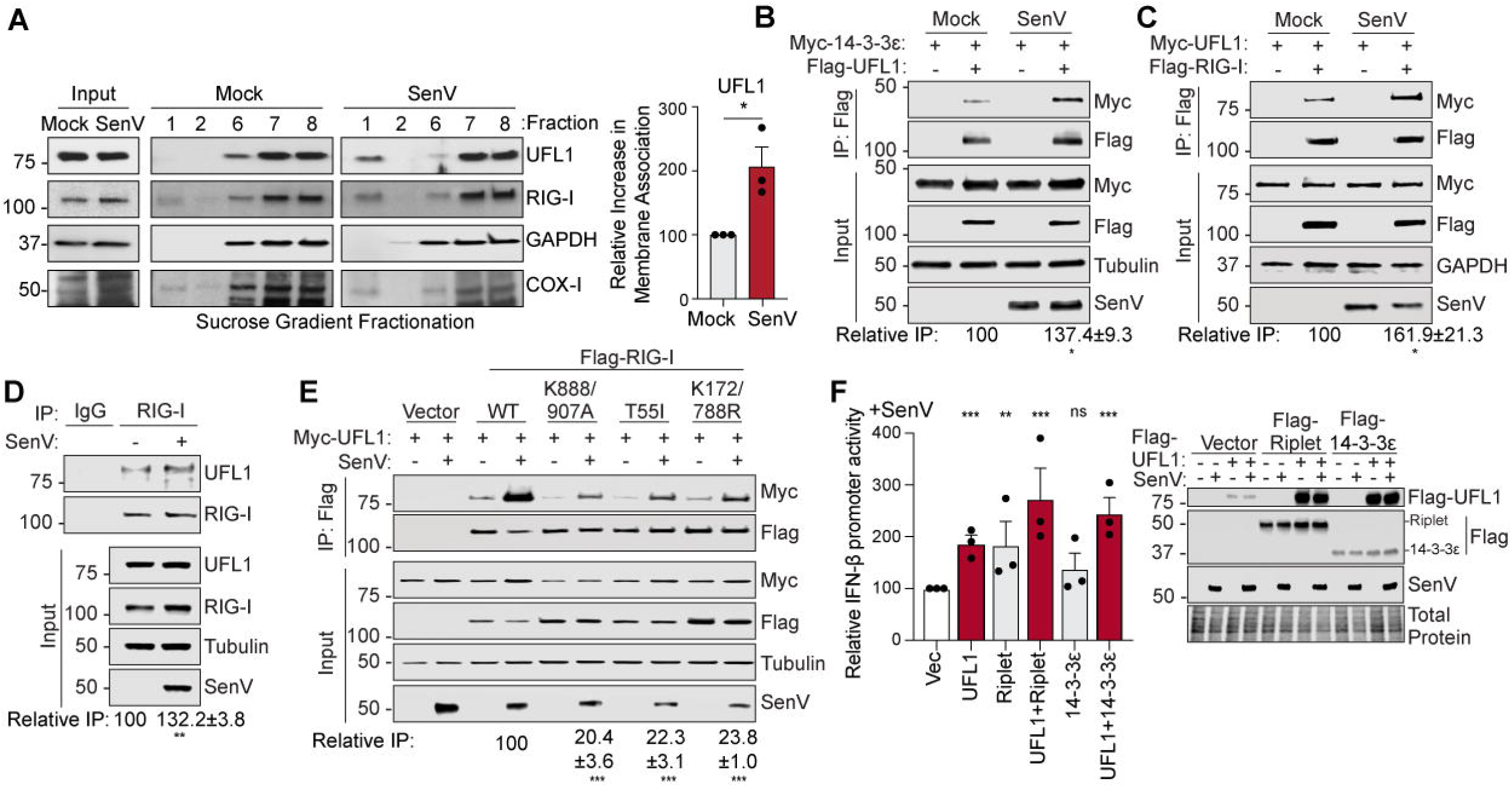
UFL1 is recruited to intracellular membranes and interacts with 14-3-3ε and RIG-I during RNA virus infection. A) Immunoblot analysis of inputs and subcellular membrane flotation of 293T cell extracts that were mock or SenV-infected (4 h) followed by sucrose gradient fractionation, with fraction numbers indicated from the top of the gradient (1) to bottom (8). Fractionation controls, GAPDH for cytosol and Cox-I for membranes, are indicated and reveal that the membranes are localized to fraction #1. Relative quantification of the ratio of UFL1 to a membrane marker (Cox-I) in fraction 1 normalized to total protein levels in inputs are shown on the right. B) Immunoblot analysis of anti-Flag immunoprecipitated extracts and inputs from 293T cells expressing Myc-14-3-3ε and Flag-UFL1 that were mock-or SenV-infected (4 h), with relative quantification on right. C) Immunoblot analysis of anti-Flag immunoprecipitated extracts and inputs from 293T cells expressing Myc-UFL1 and Flag-RIG-I that were mock-or SenV-infected (4 h), with relative quantification with IP values normalized to inputs values on right. D) Immunoblot analysis of anti-RIG-I immunoprecipitated (or anti-IgG) extracts and inputs from 293T cells that were mock- or SenV-infected (4 h), with relative quantification with IP values normalized to inputs values on right. E) Immunoblot analysis of anti-Flag immunoprecipitated extracts and inputs from 293T cells expressing Myc-UFL1 and Flag-RIG-I constructs that were mock- or SenV-infected (4 h), with results quantified as relative fold change (SenV to Mock) for each. F) Relative IFN-β-promoter reporter luciferase expression (rel. to CMV-*Renilla*) from 293T cells expressing indicated constructs followed by mock or SenV infection (18 h), with results graphed as relative SenV fold change for each. The graphs are represented as the mean -/+ SEM, n=3 (A-B, D-F) or n=4 (C) biological replicates and *p ≤ 0.05, **p ≤ 0.01, and ***p ≤ 0.001 determined by Student’s t-test (A-D) or one-way ANOVA followed by Dunnett’s multiple comparisons test (E-F).

### UFL1 interaction with RIG-I requires 14-3-3ε and UFM1

Having determined that UFL1 interacts with both activated RIG-I and 14-3-3ε following RNA virus infection, we next defined the dynamics of this complex formation by testing two distinct models. In the first model, UFL1 would interact first with activated RIG-I, induce its ufmylation, and then the UFL1-RIG-I complex would interact with 14-3-3ε. In this model, depletion of 14-3-3ε or loss of UFM1 would not be expected to change the interaction of UFL1 with RIG-I. In the second model, UFL1 would interact first with 14-3-3ε and induces its ufmylation, or that of another associated protein, and then the UFL1-14-3-3ε complex would interact with activated RIG-I. In this second model, depletion of 14-3-3ε would be expected to prevent UFL1 interaction with RIG-I, and loss of ufmylation would limit UFL1 interaction with RIG-I but would not affect UFL1 interaction with 14-3-3ε. To elucidate these possibilities, first, we used co-immunoprecipitation to measure the interaction of exogenously expressed Flag-UFL1 and HA-RIG-I in SenV-infected 293T lysates that had been depleted of 14-3-3ε or CTRL by siRNA. This revealed that formation of the SenV-activated RIG-I-UFL1 complex requires 14-3-3ε (Figure 5A). Next, we tested if ufmylation was required for formation of the SenV-activated RIG-I-UFL1 complex by measuring this interaction in WT or UFM1 KO 293T cells. We found that UFM1 was required for SenV-activated RIG-I-UFL1 complex (Figure 5B). The results of these two experiments reveal that both 14-3-3ε and UFM1 are required for UFL1 to interact with RIG-I, supporting the second model of complex formation in which UFL1 interacts first with 14-3-3ε and catalyzes its ufmylation, and then this complex associates with RIG-I. In support of this, we found that UFM1 was not required for UFL1 to interact with 14-3-3ε (Figure 5C). To test if 14-3-3ε is UFM1-conjugated, we performed an immunoprecipitation of Flag-14-3-3ε from cell extracts that were mock or SenV-infected and expressed either HA-UFM1-WT or HA-UFM1DC3, which lacks the terminal 3 amino acids required conjugation to target proteins (17). Importantly, these extracts were boiled prior to immunoprecipitation to remove non-covalent interactions. Following immunoprecipitation of Flag-14-3-3ε, immunoblotting with an anti-HA antibody revealed a slower migrating form of 14-3-3ε approximately 15 kDa heavier (HA+UFM1) than the predicted molecular weight of 14-3-3ε (37 kDa) (Figure 5D), suggestive of covalent UFM1 modification. Additionally, the proportion of 14-3-3ε, conjugated by UFM1 increases following SenV infection. Together, these data indicate that ufmylation promotes the interaction of UFL1 with 14-3-3ε and activated RIG-I and that UFM1 has increased conjugation to 14-3-3ε following RIG-I activation.

**Figure 5.**
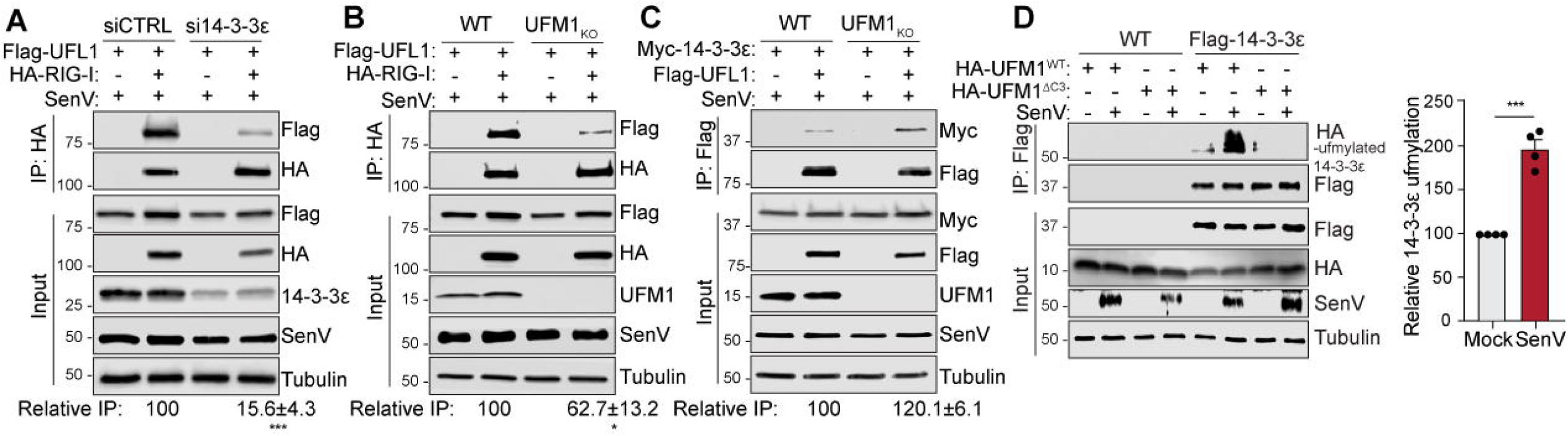
UFL1 interaction with RIG-I requires 14-3-3ε and ufmylation. A) Immunoblot analysis of anti-HA immunoprecipitated extracts and inputs from 293T cells transfected with siCTRL or si14-3-3ε followed by SenV infection (4h). B) Immunoblot of anti-HA immunoprecipitated extracts and inputs from 293T WT or UFM1 KO cells transfected with HA-RIG-I and Flag-UFL1. C) Immunoblot of anti-Flag immunoprecipitated extracts and inputs from 293T WT or UFM1 KO cells transfected with Flag-UFL1 and Myc-14-3-3ε. D) Immunoblot of anti-flag immunoprecipitated extracts and inputs from either 293T-WT or 293T-Flag-14-3-3ε cells transfected with either HA-UFM1-WT or HA-UFM1DC3. In (A-D), SenV infection was for 4 hours, and relative quantification of the interacting protein vs. IP protein is shown underneath (A-C) or as a graph (D), indicating the mean -/+ SEM (A, B, D), n=3 (A, B) or n=4 (D) biological replicates. For (C) values shown are SD of IP values adjusted for input expression, with n=2 biological replicates. *p ≤ 0.05, **p ≤ 0.01, and ***p ≤ 0.001 determined by Student’s t-test.

### Ufmylation promotes RIG-I interaction with 14-3-3ε for MAVS activation

Having found that that UFL1 requires 14-3-3ε to interact with activated RIG-I, we next tested if UFL1 is required for the interaction of 14-3-3ε with RIG-I, which is essential for activated RIG-I to translocate from the cytosol to intracellular membranes for interaction with MAVS (7). We performed a co-immunoprecipitation of Flag-RIG-I and Myc-14-3-3ε from 293T cells and found that this SenV-mediated interaction was significantly decreased upon UFL1 depletion (Figure 6A). In addition, loss of UFM1 expression also decreased the SenV-induced interaction of RIG-I with 14-3-3ε (Figure S3). Importantly, we also found that UFM1 is required for the SenV-induced interaction of RIG-I with MAVS (Figure 6B) and MAVS higher-order oligomerization, which is a hallmark of MAVS activation (45, 46) (Figure 6C). In summary, these data reveal that UFL1 and UFM1 are required for the RIG-I interaction with 14-3-3ε, for interaction with MAVS, and for MAVS activation by oligomerization.

**Figure 6.**
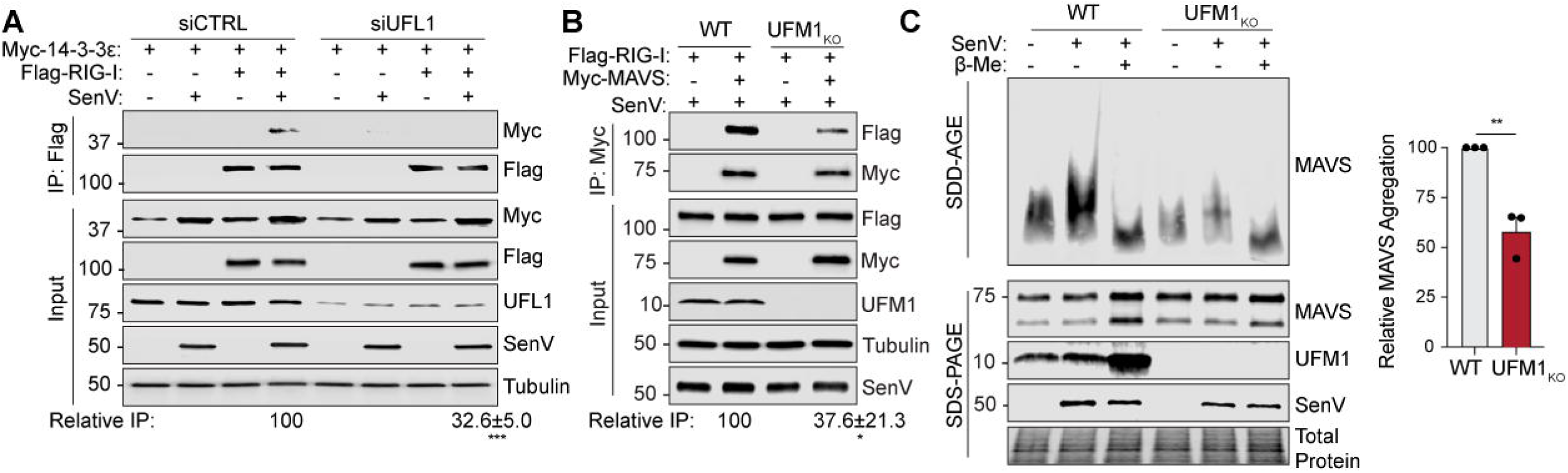
Ufmylation promotes RIG-I interaction with 14-3-3ε for MAVS activation. A) Immunoblot of anti-Flag immunoprecipitated extracts and inputs from 293T cells transfected with siCTRL or siUFL1 and indicated constructs. B) Immunoblot of anti-Myc immunoprecipitated extracts from 293T WT or UFM1 KO cells. D) 293T WT or UFM1 KO were mock or SenV-infected (12 h). Immunoblotting shows endogenous MAVS in input samples and MAVS aggregation from P5 fractions, in the presence or absence of denaturing reagent (ß-mercaptoethanol). SenV infection was for 4 h (A-C) or 12 h (D). In (A-B), relative quantification of interacting protein vs. IP protein in the IP is shown underneath; in (D) SDD-AGE MAVS values are normalized to corresponding SDS-PAGE values and shown as the ratio of SenV/mock values. Values (A-B) or graph (C) show the mean -/+ SEM for n=3 biological replicates. *p ≤ 0.05, **p ≤ 0.01, and ***p ≤ 0.001 determined by Student’s t-test.

## Discussion

Regulation of RIG-I activation and downstream signaling is essential for proper induction and termination of IFN. Here, we show that both UFL1 and the process of ufmylation promote RIG-I pathway signaling that leads to IFN induction, uncovering an important step in the activation of the RIG-I pathway. RIG-I activation occurs upon RNA binding. Then, RIG-I undergoes ATP hydrolysis. And interaction with K63-linked polyubiquitin chains, both covalently and non-covalently (5, 44, 47), which promotes formation of a RIG-I tetramer (48). This polyubiquitinated, activated RIG-I oligomer then interacts with the membrane trafficking protein 14-3-3ε for translocation to MAVS at ER-mitochondrial contact sites (7). We found that UFL1 is recruited to 14-3-3ε following RNA virus infection and that ufmylation facilitates the interaction between 14-3-3ε and activated RIG-I. Importantly, this results in increased interaction of RIG-I with MAVS and MAVS oligomerization, ultimately promoting the downstream signal transduction which produces IFN.

Ufmylation is emerging as a post-translational modification that regulates diverse biological processes, including DNA repair, ER homeostasis, and even the replication of hepatitis A virus (21, 22, 24, 27, 28, 30, 35, 49). In these cases, UFL1, along with the other members of the ufmylation cascade, induce ufmylation of a target protein important for regulating these processes. For example, both MRE11 and histone H4 are ufmylated by UFL1 in the nucleus in response to DNA damage resulting in activation the key DNA repair kinase ATM (22, 24). UFL1 can also act at the ER, where it plays a role in ER protein quality control, where it ufmylates specific proteins, including ribosomal proteins RPL26, to induce lysosomal degradation of stalled peptides and/or the ER and prevent the unfolded protein response (27, 28, 49, 50). Hepatitis A virus translation, which occurs in association with the ER, also requires ufmylation of RPL26 (30). Therefore, ufmylation can regulate several aspects of translation. It is possible that ufmylation regulates translation of certain mRNAs important for RIG-I signaling and subsequent IFN induction. However, we identified a role for ufmylation in regulating the interaction of RIG-I with 14-3-3ε, one of the earliest known steps of RIG-I signaling, strongly supporting a mechanism in which following RIG-I activation, ufmylation is controlling this specific protein-protein interaction. The mechanisms by which the process of UFM1 addition regulates interactions between proteins or alters other aspects of protein function are largely unknown. Indeed, we found that in both overexpression and KO conditions UFSP2, the protease that removes UFM1 from proteins (Figure 2D) (18), also promoted SenV-mediated IFN induction (Figure 2E-F), suggesting that we do not yet have a full grasp on the ufmylation process. It is possible that the dynamic process of ufmylation or the enhanced formation of mature UFM1 following deconjugation from targets promote RIG-I signaling independent of deconjugation activity. Interestingly, in the UFSP2 KO cells, we observed increased amounts of a specific higher molecular weight UFM1-conjugates by immunoblot, suggesting that UFSP2 is important for the generation of free UFM1 for conjugation (Figure 2F). Indeed, in support of this idea, others have shown that UFSP2 in myeloid cells is required for influenza virus resistance in mice (31). It is also possible that UFSP2 acts on other members of the RIG-I pathway to alter their function. Future studies to define how the process of ufmylation regulates this and other aspects of the antiviral innate immune response will be of great interest as they may shed light broadly on how ufmylation regulates diverse cell biological processes that alter cellular signaling.

The mechanisms underlying how cytoplasmic UFL1 is recruited to its protein targets that reside in different subcellular compartments are not fully known. For example, we found that RIG-I activation induces UFL1 translocation to intracellular membranes (Figure 3A), and while we know that UFL1 is recruited to the MAM during infection, the mechanism by which UFL1 becomes membrane-associated remains unknown (14). DDRGK1 (UFBP1) may facilitate UFL1 targeting to the MAM, as DDRGK1 is localized to mitochondrial-ER contact sites (16, 51) and in some cases it is required for UFL1 recruitment to membranes (27, 28). Thus, both DDRGK1 and mitochondrial-ER contact sites could function as a regulatory hub that aids in the recruitment of UFL1 and RIG-I pathway signaling proteins. Interestingly, RAB1B, a GTPase that we found is recruited to the MAM and important for RIG-I signaling (14, 15) is ufmylated (52, 53), which reveals that ufmylation likely regulates a number of RIG-I pathway signaling proteins. As UFL1 contains no functional domains common to other E3 ligases that might allow one to predict how its targets are selected (16, 54, 55), defining the signals and features that control UFL1 localization, as well as the target proteins ufmylated in response to RIG-I activation, will undoubtedly reveal clues into how the process of ufmylation is activated and how specific targets are selected.

Our work revealed that 14-3-3ε required ufmylation to interact with activated RIG-I. The details underlying how 14-3-3ε interacts with activated RIG-I have not been fully elucidated, as it does not occur through the known phosphorylated amino acids on RIG-I, the typical recruitment signal of the 14-3-3 family of proteins (7, 56, 57). Our work supports the work of others demonstrating that 14-3-3ε interacts with UFM1 and other members of the ufmylation pathway, and it reveals that SenV increases the proportion of ufmylated 14-3-3ε (Figure 5D) (16). It is likely that 14-3-3ε is mono-ufmylated as the apparent molecular weight of 14-3-3ε increased by one ufmylation group (HA+UFM1, 15 kDa; Figure 5D). Thus, taken together with our results, this suggests that ufmylation of 14-3-3ε or a 14-3-3ε-associated protein promotes the interaction between activated RIG-I and 14-3-3ε. In fact, a number of 14-3-3 family proteins are post-translationally modified by phosphorylation, acetylation, and oxidation (58). Therefore, post-translational modification of 14-3-3ε by ufmylation could define how cargo proteins, including RIG-I, are selected. Indeed, this mechanism could be shared with other RNA virus sensing pathways, such as the RIG-I-like-receptor MDA5, which also interacts with a 14-3-3 protein, 14-3-3η, by an unknown mechanism (59). Thus, ufmylation may broadly influence how 14-3-3 proteins or other host proteins interact with each other to regulate the intracellular innate immune response.

Overall, this work lays the groundwork for future studies to define how ufmylation of antiviral innate immune signaling proteins regulates their function and how specific signaling pathways are differentially activated through ufmylation. In addition, our work adds ufmylation to the growing list of ubiquitin-like and other modifications that regulate the intracellular innate immune response, including ISGylation, SUMOylation, FATylation, acetylation, phosphorylation, and others (4, 60, 61) broadening our understanding of how RIG-I signaling is activated and rapidly controlled by post-translational modifications in response to infection, leading to greater knowledge of the exquisite regulation of these pathways.

## Materials and Methods

### Cell lines, viruses, and treatments

Neonatal human dermal fibroblast (NHDF) cells and embryonic kidney 293T cells were grown in Dulbecco’s modification of Eagle’s medium (DMEM; Mediatech) supplemented with 10% fetal bovine serum (Thermo Fisher Scientific), 1X minimum essential medium non-essential amino acids (Thermo Fisher Scientific), and 25 mM HEPES (Thermo Fisher Scientific) (cDMEM). 293T (CRL-3216) were obtained from American Type Culture Collection (ATCC), NHDF cells (CC-2509) were obtained from Lonza. All cell lines were verified as mycoplasma free by the LookOut Mycoplasma PCR detection kit (Sigma). SenV Cantell strain was obtained from Charles River Laboratories and used at 200 hemagglutination units/mL (HAU). SenV infections were performed in serum-free media (30 minutes to 1 hour), after which complete media was replenished. IFN-β (PBL Assay Science) was added to cells at a concentration of 50 units/mL in cDMEM for 18 hours.

### Plasmids

The following plasmids have been previously described: pEF-TAK-Flag, pEF-BOS-Flag-RIG-I (62), pEF-TAK-Flag-Riplet (63) pIFN-β-luc (64), pCMV-Renilla (Promega), pX459 (Addgene #62988), psPAX2 (Addgene #12260), and pMD2.G (Addgene #12259), pEF-BOS-Flag-RIG-I T55I (65), pEF-TAK-Myc-MAVS (32). pLJM1_Flag-UFM1 was a gift from Drs. Craig McCormick and John Rohde at Dalhouise University. The following plasmids were generated by insertion of PCR-amplified fragments into the NotI-to-PmeI digested pEF-TAK-Flag using InFusion cloning (Clontech): pEF-TAK-Flag-UFL1 (GenBank: BC036379; GeneID: 23376), pEF-TAK-Flag-UBA5 (NM_024818.6), pEF-TAK-Flag-UFC1 (NM_016406.4), pEF-TAK-Flag-UFSP2 (NM_018359.5), pEF-TAK-Flag-UFL1 1-212, pEF-TAK-Flag-UFL1 1-452, pEF-TAK-Flag-UFL1 213-794, pEF-TAK-Flag-UFL1 453-794, and pEF-TAK-Flag-14-3-3ε. Both pEF-TAK-Myc-14-3-3e and pEF-TAK-Myc-UFL1 were generated by insertion of PCR-amplified fragments into the AgeI-NotI digested pEF-TAK-Myc (pEF-TAK-Myc-MAVS) by InFusion. The pEF-TAK-HA vector was generated by PCR to replace Flag with HA, and pEF-TAK-HA-RIG-I, and pEF-TAK-HA-UFM1 were generated by insertion of a PCR-amplified fragment into the NotI-AgeI digested pEF-TAK-HA vector. pLEX-Flag-14-3-3e was generated by ligation of a PCR-amplified Flag-14-3-3e into the BamHI-to-XhoI digested pLEX using InFusion cloning. The following plasmids were generated by site-directed mutagenesis: pEF-TAK-Flag-UFL1^siR^, pEF-TAK-HA-UFM1DC3, pEF-BOS-Flag-RIG-I K888/907A, and pEF-BOS-Flag-RIG-I K172/788R. To generate the CRISPR guide RNA plasmids px459-UFM1-E2, px459-UFM1-B, and px459-UFSP2 sgRNA oligonucleotides were annealed and inserted into the BbsI-digested pX459 (30, 66). The plasmid sequences were verified by DNA sequencing and oligonucleotide sequences are available upon request.

### Generation of RNA PAMP

Annealed oligonucleotides containing the sequence of the HCV 5’ppp poly-U/UC region (34) were *in vitro* transcribed using the MEGAshortscript T7 transcription kit (Ambion) followed by ethanol precipitation, with the resulting RNA resuspended at 1 μg/μL.

### Transfection

DNA transfections were performed using FuGENE6 (Promega) or TransIT-LT1 (Mirus Bio). RNA PAMP transfections were done using the TransIT-mRNA Transfection kit (Mirus Bio). The siRNA transfections were done using Lipofectamine RNAiMax (Invitrogen). siRNAs directed against 14-3-3ε (Dharmacon-L-017302-02-0005), UFL1 (Qiagen-SI04371318) or non-targeting AllStars negative control siRNA (Qiagen-1027280) were transfected into 293T cells (25 pmol of siRNA; final concentration of 0.0125 μM) or NHDF cells (250 pmol of siRNA; final concentration of 0.25 μM). Media was changed 4-24 hours post-transfection, and cells were incubated for 36-48 h post-transfection prior to each experimental treatment. IFN-β-promoter luciferase assays were performed as previously described at 18-24 hours post treatment and normalized to the *Renilla* luciferase transfection control (33).

### ELISA

IFN-β ELISAs were performed using Human IFN-beta DuoSet (R&D Systems) with supernatants collected from cultured cells.

### Generation of cell lines

UFM1 and UFSP2 KO 293T cells were generated by CRISPR/Cas9, using two guides targeting exon 2 and 3 (UFM1) or a single guide targeting exon 6 (UFSP2), similar to others, as we have done previously. Single cell clones were validated via anti-UFM1 or anti-UFSP2 immunoblot and genomic sequencing, with one clone used here (15, 30). 293T cell pools overexpressing Flag-14-3-3e were generated by lentiviral transduction, as previously (36).

### RNA analysis

Total cellular RNA was extracted using the RNeasy Plus mini kit (Qiagen). RNA was then reverse transcribed using the iScript cDNA synthesis kit (BioRad), as per the manufacturer’s instructions. The resulting cDNA was diluted 1:3 in ddH2O. RT-qPCR was performed in triplicate using the Power SYBR Green PCR master mix (Thermo-Fisher) and QuantStudio 6 Flex RT-PCR system. Oligonucleotide sequences for qPCR are available upon request.

### RNA-seq

WT and UFM1 KO 293T cells were mock or SenV-infected (18 h) and harvested in biological duplicate, followed by total RNA extraction via TRIzol reagent (Thermo Fisher Scientific). Sequencing libraries were prepared using the KAPA Stranded mRNA-Seq Kit (Roche) and sequenced on an Illumina Novaseq 6000 with 50 bp paired-end reads (>20 million reads per sample) in an S1 flow cell by the Duke University Center for Genomic and Computational Biology. RNA-seq data was processed using the TrimGalore toolkit (67) which employs Cutadapt (68) to trim low-quality bases and Illumina sequencing adapters from the 3’ end of the reads. Only reads that were 20nt or longer after trimming were kept for further analysis. Reads were mapped to the GRCh38v93 version of the human genome and transcriptome (69) using the STAR RNA-seq alignment tool (70). Reads were kept for subsequent analysis if they mapped to a single genomic location. Gene counts were compiled using the HTSeq tool (71). Only genes that had at least 10 reads in any given library were used in subsequent analysis. Normalization and differential expression was carried out using the DESeq2 (72) Bioconductor (73) package with the R statistical programming environment. The false discovery rate was calculated to control for multiple hypothesis testing. Gene set enrichment analysis (74) was performed to identify gene ontology terms and pathways associated with altered gene expression for each of the comparisons performed. All RNA-seq data are deposited in the GEO database under GSE186287.

### Immunoblotting

Cells were lysed in a modified radioimmunoprecipitation assay (RIPA) buffer (10 mM Tris [pH 7.5], 150 mM NaCl, 0.5% sodium deoxycholate, and 1% Triton X-100) supplemented with protease inhibitor cocktail (Sigma) and Halt Phosphatase Inhibitor (Thermo-Fisher), and post-nuclear lysates were isolated by centrifugation. Quantified protein (between 5 −15 μg) was resolved by SDS/PAGE, transferred to nitrocellulose or polyvinylidene difluoride (PVDF) membranes in a 25 mM Tris-192 mM glycine-0.01% SDS buffer. Membranes were stained with Revert 700 total protein stain (LI-COR Biosciences) and then blocked in 3% BSA in Tris-buffered saline containing 0.01% Tween-20 (TBS-T). After washing with PBS-T or TBS-T (for phosphoproteins) buffer, following incubation with primary antibodies, membranes were incubated with species-specific horseradish peroxidase-conjugated antibodies (Jackson ImmunoResearch, 1:5000) or fluorescent secondaries (LI-COR Biosciences), followed by treatment of the membrane with Clarity Western ECL substrate (BioRad) and imaging on a LICOR Odyssey FC. The following antibodies were used for immunoblotting: R-anti-SenV (MBL, 1:1000), M-anti-Tubulin (Sigma, 1:1000), R-anti-GAPDH (Cell Signaling Technology, 1:1000), R-anti-p-IRF3 (Cell Signaling Technology, 1:1000), R-anti-IRF3 (Cell Signaling Technology, 1:1000), R-anti-UFL1 (Novus Biologicals, 1:1000), R-anti-UFM1 (Abcam, 1:1000), anti-RIG-I (M-AdipoGen, R-Abcam, 1:1000), R-anti-14-3-3ε (Cell Signaling Technology, 1:1000), M-anti-Flag M2 (Sigma, 1:1000), anti-Flag-HRP (Sigma, 1:1000-1:5000), R-anti-Flag (Sigma, 1:1000), anti-HA (M- and R-Sigma, 1:1000), and anti-Myc (M-Santa Cruz or R-Cell Signaling Technology, 1:1000).

### Immunoprecipitation

Cells were lysed in RIPA buffer with or without 10% glycerol. Quantified protein (between 100-500 μg) was incubated with protein-specific, isotype control antibody (R-Cell Signaling Technology or M-Thermo Fisher), or anti-Flag M2 magnetic beads (Sigma), in lysis buffer either at room temperature for 2 h or at 4°C overnight with head over tail rotation. The lysate/antibody mixture was then incubated with Protein G Dynabeads (Invitrogen) for 1 h. Beads were washed 3X in PBS or RIPA buffer and eluted in 2X Laemmli Buffer (BioRad) with or without 5% 2-Mercaptoethanol at 95°C for 5 min. Proteins were resolved by SDS/PAGE and immunoblotting, as above. For ufmylation immunoprecipitations, cells were lysed with NP-40 buffer (50 mM Tris [pH 8], 150 mM NaCl, 0.5% sodium deoxycholate, and 1% NP-40) supplemented with 10% glycerol, protease and phosphatase inhibitors, as above, and N-ethylmaleimide (Sigma-Aldrich). Post-nuclear lysates were boiled at 95°C for 5 min and incubated with anti-Flag M2 magnetic beads, as above.

### Subcellular membrane fractionation

Membrane fractionation was performed as previously described (7, 12, 38, 75). Cells were lysed in hypotonic buffer (10 mM Tris-HCL 7.5), 10 mM KCl, and 5 mM MgCl_2_ supplemented with protease inhibitor cocktail) for 10 minutes on ice followed by 20 passages through a 20-guage needle. Nuclei and unbroken cells were removed by centrifugation at 1000xg for 5 min at 4°C. The resulting supernatants were mixed thoroughly with 72% sucrose and overlayed with 55% sucrose, followed by 10% sucrose, all in low-salt buffer (2 nM EDTA, 20 nM HEPES (pH 8.0), 150 mM NaCl, 0.1% SDS, 1% Triton X-100). The gradients were subjected to centrifugation at 38,000 RPM in a Beckman SW41 Ti Rotor for 14 h at 4°C. 1 mL fractions were collected using a BioComp piston gradient fractionator and resulting fractions were divided in half and mixed with 2 parts 100% methanol and precipitated overnight at −80°C. Protein pellets were collected by centrifugation and resuspended in 2X Laemmli buffer and heated for 5 min at 95°C for immunoblot analysis. 10% pre-fractionated cells from each condition were collected as the input.

### Semi-denaturing detergent agarose gel electrophoresis

SDD-AGE was performed as described (45, 46). Briefly, crude mitochondria (P5 fraction) were isolated from an equal number of WT or UFM1 KO 293T cells that were mock or SenV infected (12 h), resuspended in hypotonic buffer (10 mM Tris, pH 7.5, 10 mM KCl, 1.5 mM MgCl_2_, and 0.5 mM EDTA). Resulting samples were split and 2X SDD-AGE sample buffer (0.5X TBE, 10% glycerol, 2% SDS, 0.2 mM Bromophenol Blue) buffer with or without 5% 2-Mercaptoethanol was added, and samples were loaded onto a vertical 1.5% agarose gel. Electrophoresis was performed with a constant voltage of 70 V at 4 °C in SDD-AGE running buffer (1X TBE and 0.1% SDS). Gels were transferred onto a nitrocellulose membrane overnight on ice at 25 V. Membranes were fixed in 0.25% glutaraldehyde in PBS and immunoblotting was performed as usual. 15% of the SDD-AGE samples were reserved for input.

### Quantification of immunoblots

Immunoblots imaged using the LICOR Odyssey FC were quantified by ImageStudio software, and raw values were normalized to relevant controls for each antibody. Phosphoprotein values were normalized to Tubulin and displayed as the percentage of signal from WT. Relative membrane association of UFL1 was quantified as the ratio of UFL1 to Cox-1 in fraction 1 normalized to total protein levels of UFL1 in the input and displayed as the percentage of UFL1 membrane association normalized to mock values.

### Statistical analysis

Student’s unpaired t-test, one-way ANOVA, or two-way ANOVA were implemented for statistical analysis of the data followed by appropriate post-hoc test (as indicated) using GraphPad Prism software. Graphed values are presented as mean ± SD or SEM (n = 3 or as indicated); *p ≤ 0.05, **p ≤ 0.01, and ***p ≤ 0.001.

## Acknowledgments

We thank colleagues who provided reagents (see Methods), the Duke Functional Genomics Core Facility, the Duke University Center for Genomic and Computational Biology, the Duke Genomic Analysis and Bioinformatics Shared Resource and Dr. Wei Chen for performing RNA-seq analysis, Dr. Madhuvanthi Vijayan and Boyoung Michelle Kim for generation and initial characterization of plasmids, Dr. Matthew Thompson for input in RNA-seq sample preparation and data processing, and Horner lab members for useful discussion. This work was supported by funds from Burroughs Wellcome Fund, National Institutes of Health: R21AI144380, R01AI155512, T32-CA009111, and an American Cancer Society Postdoctoral fellowship 131321-PF-17-188-01-MPC.

## Author Contributions

Conceptualization: D.L.S., D.C.B., and S.M.H.; Investigation: D.L.S, D.C.B., and M.P.; Formal analysis: D.L.S., D.C.B., K.A.M., and S.M.H.; Writing – original draft: D.L.S. and S.M.H.; Writing – review and editing: D.C.B., D.L.S., and S.M.H. All authors read the manuscript and provided comments. Funding acquisition: S.M.H.

---

**Figure S1.**
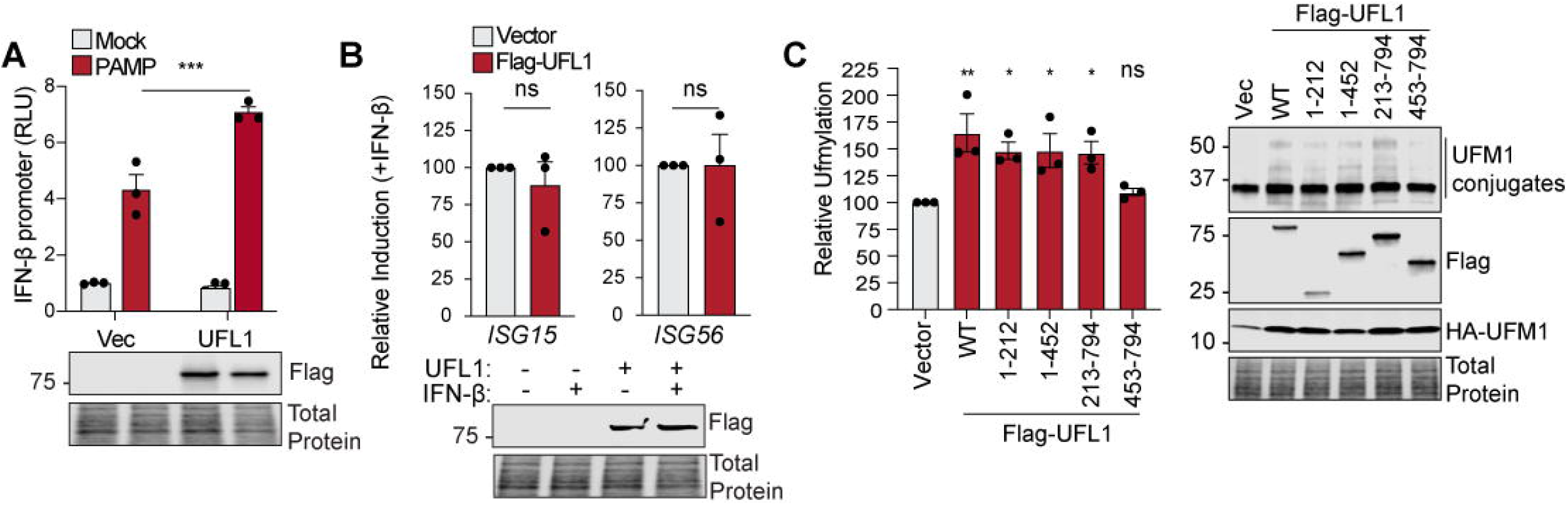
UFL1 positively regulates IFN induction. A) IFN-β-promoter reporter luciferase expression (rel. to CMV-*Renilla*) from 293T cells transfected with vector (Vec) or Flag-UFL1, followed by mock or HCV PAMP RNA transfection (24 h). B) RT-qPCR analysis (rel. to *GAPDH*) of RNA extracted from 293T cells transfected vector or Flag-UFL1 that were treated with IFN-β (18 h). C) Quantification of immunoblots from 293T cells expressing HA-UFM1, and either vector or indicated Flag-UFL1 represented as the ratio of UFM1 conjugates (approximately 25-50 kDa) to total protein expression in each lane with vector set to 100. Graph indicates the mean -/+ SEM for n=3 biological replicates. *p ≤ 0.05 and **p ≤ 0.01 as determined by two-way ANOVA followed by Šidák’s multiple comparisons test (A), Student’s t-test (B) or one-way ANOVA followed by Dunnett’s multiple comparisons test (C).

**Figure S2.**
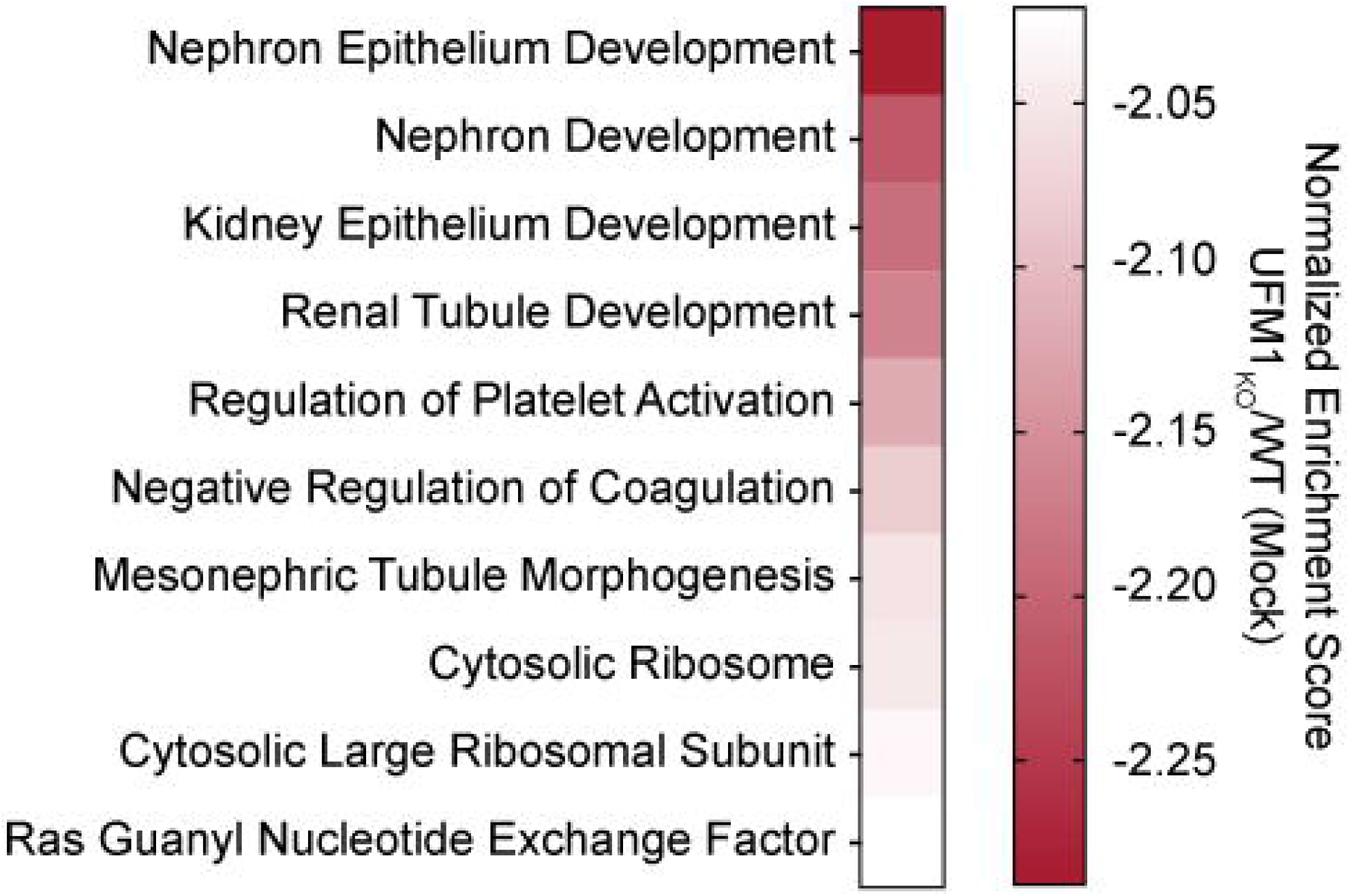
Transcriptional response of genes negatively regulated by UFM1. RNA-seq analysis of WT versus UFM1 KO 293T cells showing the gene set enrichment analysis (top 10 categories) of negatively regulated differentially expressed genes represented by normalized enrichment score to identify gene ontology terms and pathways associated with altered gene expression for each of the comparisons performed (adj P<0.01).

**Figure S3.**
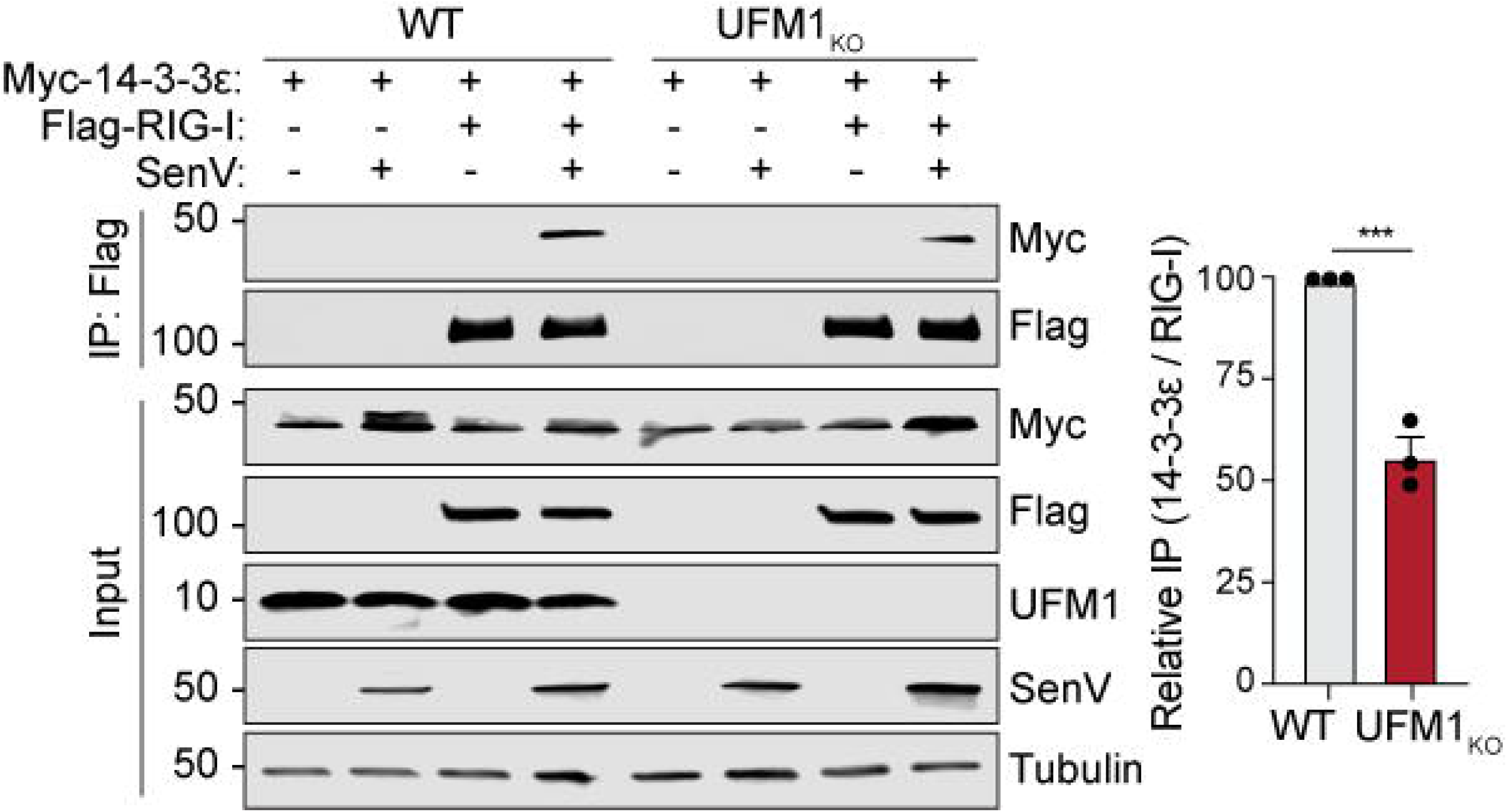
UFM1 is required for the interaction of RIG-I and 14-3-3ε. Immunoblot of anti-Flag immunoprecipitated extracts and inputs from 293T WT or UFM1 KO cells with relative quantification on the right. Graph shows the mean -/+ SEM for n=3 biological replicates; ***p ≤ 0.001 determined by Student’s t-test.

**Dataset S1. Differential expression analysis from RNA-seq analysis for UFM1 KO / WT 293T cells**

Table S1.1: UFM1 KO / WT Mock

Table S1.2 UFM1 KO / WT SenV (18 h)

**Dataset S2. Gene Set Enrichment Analysis for UFM1 KO / WT 293T cells**

Table S2.1: UFM1 KO / WT Mock- negative direction

Table S2.2 UFM1 KO / WT SenV (18 h)- negative direction

Table S2.3 UFM1 KO / WT Mock- positive direction

Table S2.4 UFM1 KO / WT SenV (18 h)- negative direction

